# Regulation of Hippocampal Excitatory Synapses by the Zdhhc5 Palmitoyl Acyl Transferase

**DOI:** 10.1101/2020.09.11.294397

**Authors:** Jordan J. Shimell, Andrea Globa, Marja D. Sepers, Angela R. Wild, Nusrat Matin, Lynn A. Raymond, Shernaz X. Bamji

**Author notes:** Co-first authorship (Jordan J. Shimell, Andrea Globa). CORRESPONDING AUTHOR: Shernaz X. Bamji, Ph.D., Department of Cellular & Physiological Sciences, University of British Columbia, 2350 Health Sciences Mall, Vancouver, B.C., V6T 1Z3, Canada.

## Abstract

Palmitoylation is the most common post-translational lipid modification in the brain; however, the role of palmitoylation and palmitoylating enzymes in the nervous system remains elusive. One of these enzymes, Zdhhc5, has previously been shown to regulate synapse plasticity. Here, we report that Zdhhc5 is also essential for the formation of excitatory, but not inhibitory synapses both *in vitro* and *in vivo*. We demonstrate *in vitro* that this is dependent on Zdhhc5’s enzymatic activity, its localization at the plasma membrane, and its C-terminal domain which has been shown to be truncated in a patient with schizophrenia. Loss of Zdhhc5 in mice results in a decrease in the density of excitatory hippocampal synapses accompanied by alterations in membrane capacitance and synaptic currents, consistent with an overall decrease in spine number and silent synapses. These findings reveal an important role for Zdhhc5 in the formation and/or maintenance of excitatory synapses.

## INTRODUCTION

Palmitoylation is a reversible post-translational lipid modification that anchors proteins to specialized membrane domains and can critically impact protein stability, trafficking and function (Aicart-Ramos et al., 2011; Hannoush & Sun, 2010; Linder & Deschenes, 2007; Resh, 2006; Salaun et al., 2010). Palmitoylation is mediated by a family of 24 ZDHHC enzymes (nomenclature in humans; Braschi et al., 2019), and growing evidence suggests that ZDHHC enzymes are important for proper brain development and function. Indeed, palmitoylation is the most common lipid modification in the brain (Fukata & Fukata, 2010) and has been shown to regulate numerous neuronal processes, including neurite outgrowth, axon pathfinding, filopodial formation, and spine development, maintenance, pruning and plasticity (Arstikaitis et al., 2008; Gauthier-Campbell et al., 2004; El-Husseini et al., 2002; Kato et al., 2000; Kutzleb et al., 1998; Laux et al., 2000; Ueno, 2000; Brigidi et al., 2015,Thomas et al., 2012; Hayashi et al., 2005; Greaves & Chamberlain, 2011; Kang, 2004; Kang et al., 2008, Shah et al., 2019; Shimell et el., 2019). Of note, 9 of the 24 ZDHHC enzymes are associated with disorders of the brain (reviewed in Zareba-Koziol et al., 2018), and bioinformatics analysis has demonstrated that while 10% of all gene products are modified by palmitoylation, 41% of all synaptic proteins (Sanders et al., 2015) are substrates for palmitoylation, further amplifying the potential role for palmitoylation in synapse biology.

Our lab has previously shown an important role for one of these enzymes, ZDHHC5, in regulating the plasticity of synaptic connections in the hippocampus (Brigidi et al., 2015). In this study, the dynamic trafficking of ZDHHC5 enabled differential palmitoylation of its substrates, providing one means by which zDHHC5 function can be regulated. ZDHHC5 can be stabilized at the synaptic membrane through its association with its accessory protein, Golga7b (Woodley & Collins, 2019) as well as Fyn kinase (Brigidi et al., 2015), by inhibiting ZDHHC5-AP2µ interactions and clathrin-mediated endocytosis. Binding to PSD-95 via its C-terminal PDZ-binding domain further stabilizes ZDHHC5 at the membrane (Brigidi et al., 2015). Interestingly, a *de novo* mutation in ZDHHC5 has been reported in a patient with schizophrenia that introduces a premature stop codon at residue 648 (E648), resulting in the loss of the last 68 amino acids of zDHHC5 (Fromer et al., 2014), including the PDZ-binding motif (Li et al., 2010; Brigidi et al., 2015). These studies reveal that ZDHHC5 is closely associated with a number of binding partners that control its subcellular localization and function.

In the present study we demonstrate both *in vitro* and *in vivo* that Zdhhc5 (mouse nomenclature; Braschi et al., 2019) can regulate the formation and/or maintenance of excitatory, but not inhibitory, synapses. Moreover, we show that this depends on Zdhhc5’s enzymatic activity, plasma membrane localization and C-terminal domain. This, coupled with the findings that Zdhhc5 substrates are localized to different subcellular domains (Li et al., 2012; Thomas et al., 2012; Brigidi et al., 2015; Kokkola et al., 2011) and the fact that Zdhhc5 localization can be dynamically regulated (Brigidi et al., 2015) suggests that Zdhhc5 may play a key role in the regulation of excitatory synapses during development by changing its subcellular localization in response to external cues.

## RESULTS AND DISCUSSION

### Zdhhc5 promotes excitatory synapse formation and regulates spine stability

To determine whether Zdhhc5 is involved in dendritic arborization and/or the formation of excitatory and inhibitory synapses, hippocampal neurons were transfected with eGFP plus the indicated constructs and immunostained for PSD-95 and gephyrin (faithful markers of excitatory and inhibitory synapses, respectively (Brigidi et al., 2015; Shah et al., 2019; Shimell et al., 2019); masking shown in Supplementary Figure 1A, B). While Zdhhc5 knockdown did not impact dendritic length (Figure 1A), complexity (Figure 1B), or the density of gephyrin puncta (Figure 1C, D), it did significantly reduce the density of PSD-95 puncta (Figure 1C, D; see Supplementary Figure 1B for the creation of masks to quantify synaptic marker density). We confirmed that the reduction in PSD-95 puncta reflected a decrease in *bona fide* excitatory synapses by quantifying the density of colocalized PSD-95/VGlut1 puncta (Figure 1E). A reduction in the density of dendritic spines, and specifically a decrease in the density of stubby and mushroom spines was also observed in Zdhhc5 knockdown neurons (Figure 1F, G).

**Figure 1.**
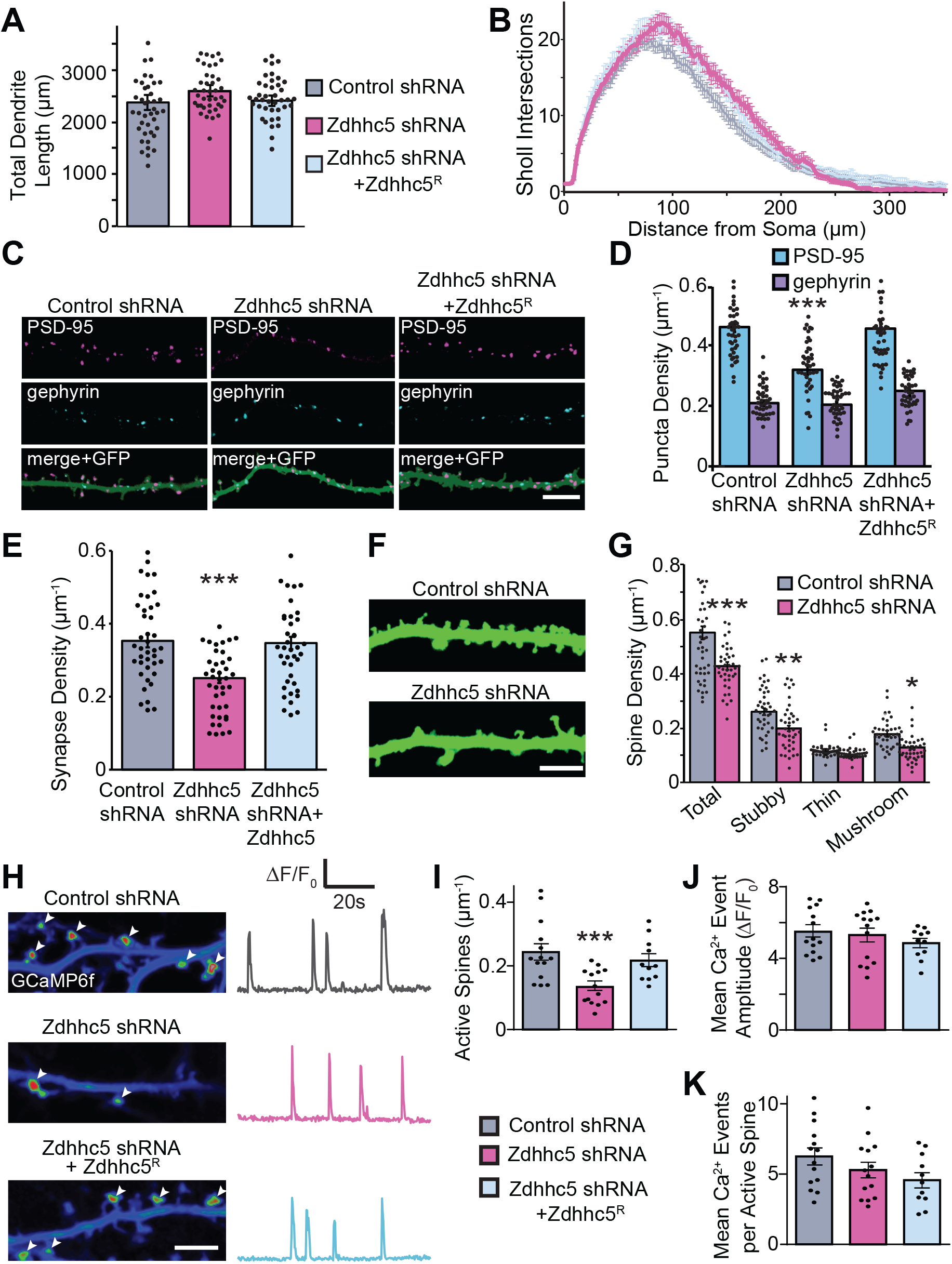
Zdhhc5 regulates the density of excitatory synapses and mature spines. (A, B) Transfection with Zdhhc5 shRNA ± WT Zdhhc5R had no effect on total dendritic length (A) or complexity (B) in cultured hippocampal neurons (13 DIV). (C) Representative images of PSD-95 and gephyrin immunolabeling with GFP cell fill. Scale bar 5 µm. (D-E) Zdhhc5 shRNA significantly reduced the density of the excitatory post-synaptic marker, PSD-95 (D) as well as colocalized PSD-95/VGlut-1 puncta (E), with no changes in the density of the inhibitory post-synaptic marker, gephyrin (D). These effects were reversed by co-transfection with Zdhhc5R. n = 40 neurons per condition, >3 cultures. (F, G) Zdhhc5 shRNA significantly reduced the density of total spines, specifically resulting from a reduction in stubby and mushroom spines. n = 40 neurons per condition, 3 cultures. (H) Left: Pseudocolored maximum intensity projected time-lapse images of GCaMP6f fluorescence from a 2 min acquisition showing the locations of miniature spine Ca2+ events (white arrows) imaged in TTX (1 μM; 15 DIV). Scale bar 5 µm. Right: GCaMP6f ΔF/F0 traces from representative single spine ROIs. (I-K) Zdhhc5 knockdown decreased the density of active spines (I), but not the amplitude (J) or number of Ca2+ events per spine (K). n = 11-14 neurons per condition, 3 cultures. *p<0.05, **p<0.01, ***p<0.001; one-way ANOVA; Tukey’s post hoc; mean ± SEM.

We next measured miniature spine Ca^2+^ transients with GCaMP6f to determine whether Zdhhc5 knockdown alters synaptic activity (Reese and Kavalali, 2015; Sinnen et al., 2016). The density of active spines was significantly reduced in cells expressing Zdhhc5 shRNA (Figure 1I). However, no changes were observed in the amplitude of spine Ca^2+^ events (Figure1 H, J) or the mean number of Ca^2+^ events at each active spine (Figure1 H, K), indicating that Zdhhc5 may regulate excitatory synapse density without altering other properties of basal synaptic transmission.

### Zdhhc5 domains important for regulating synapse density

Zdhhc5 has an extended C-terminal tail (∼70% of the total protein length) that contains a PDZ-binding motif (Kim & Sheng, 2004; Feng & Zhang, 2009; Li et al., 2010; Brigidi et al., 2015), as well as phosphorylation and palmitoylation sites that regulate internalization (Brigidi et al., 2015; Woodley & Collins, 2019). To investigate the mechanism by which Zdhhc5 regulates synapse formation and/or maintenance, we tested the ability of several Zdhhc5 mutants to rescue the Zdhhc5 knockdown-mediated decrease in excitatory synapse density. As Zdhhc5 membrane localization is activity regulated and plays an important role in synapse plasticity (Brigidi et al, 2015), we also examined the membrane localization of each mutant.

None of the Zdhhc5 mutant constructs impacted the density of gephyrin puncta (Figure 2A, B). While expression of shRNA-resistant Zdhhc5^R^ rescued the Zdhhc5 knockdown-mediated decrease in PSD-95 puncta density (Figure 1, C-E; 2A, B), expression of the enzymatically-dead mutant, Zdhhs5^R^ (in which the cysteine residue in the catalytic DHHC domain was changed to serine) did not, indicating that the enzymatic activity of Zdhhc5 is required for the formation and/or maintenance of excitatory synapses (Figure 2A-C). Surface levels of Zdhhs5^R^ were similar to wildtype Zdhhc5^R^, suggesting that the inability of Zdhhs5^R^ to rescue the Zdhhc5 knockdown phenotype was due to its lack of enzymatic function and not changes in surface expression (Figure 3A, B).

**Figure 2.**
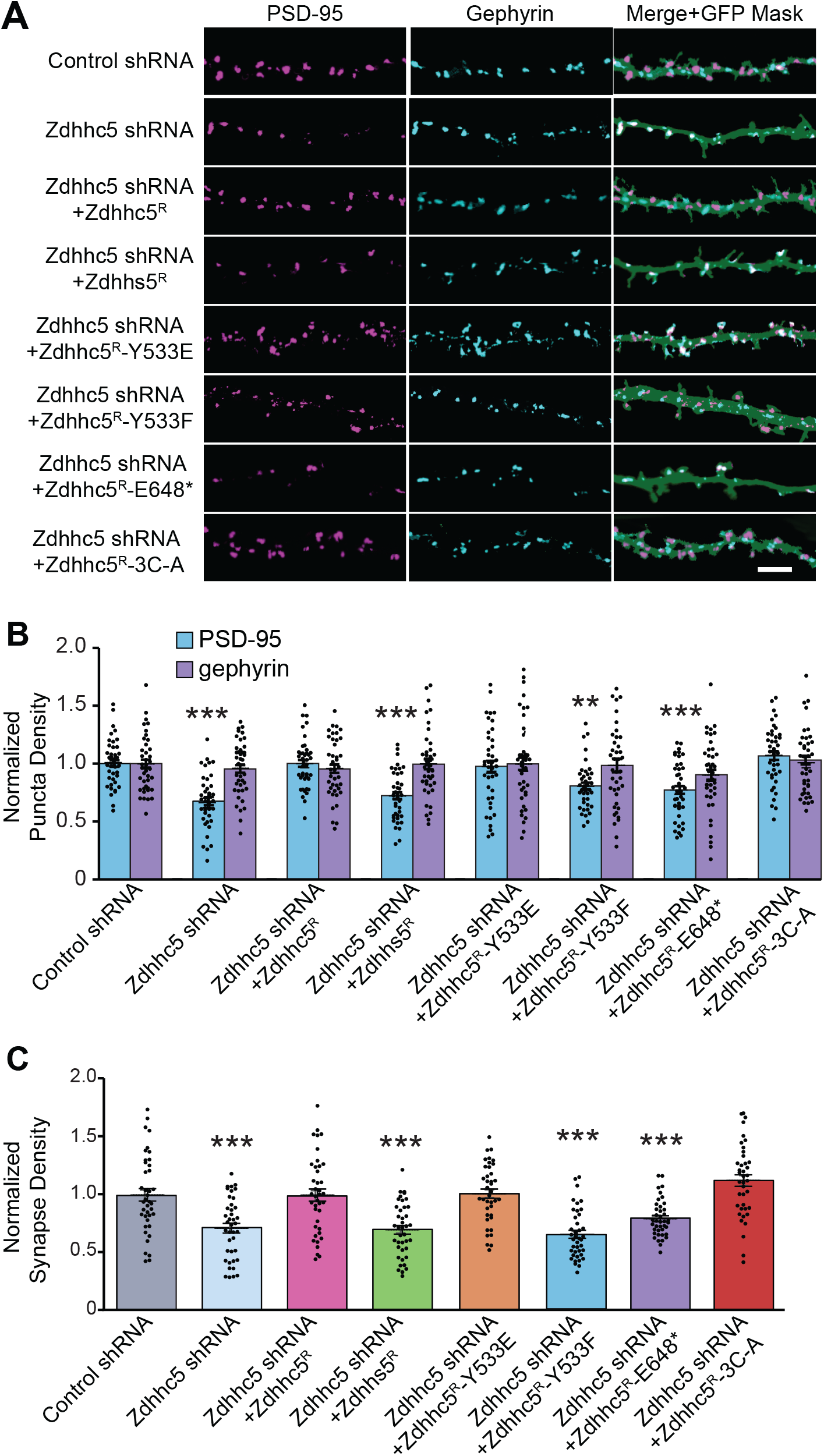
Zdhhc5 regulates excitatory synapse density via its enzymatic activity, plasma membrane localization, and C-terminal domain. (A) Representative confocal images of rat hippocampal neurons transfected at 10 DIV with eGFP plus the indicated constructs and imaged at 13 DIV. Scale bar = 5 µm. (B,C) The normalized density of PSD-95-positive puncta (B) and co-localized PSD-95/VGlut1 puncta (C) was significantly decreased in cells expressing Zdhhc5 shRNA. Synapse density was rescued by the Zdhhc5 Y533E (phospho-mimetic) and the 3C-A (palmitoylation-dead) mutants, but not the Zdhhs5 (enzymatically-dead), Zdhhc5 Y533F (phospho-dead) or E648* (C-terminal truncation) mutants. The density of gephyrin-positive puncta remained unchanged. n = 40 neurons per condition, >3 cultures. **p < 0.01, ***p < 0.001; one-way ANOVA; Tukey’s post hoc; mean ± SEM.

**Figure 3.**
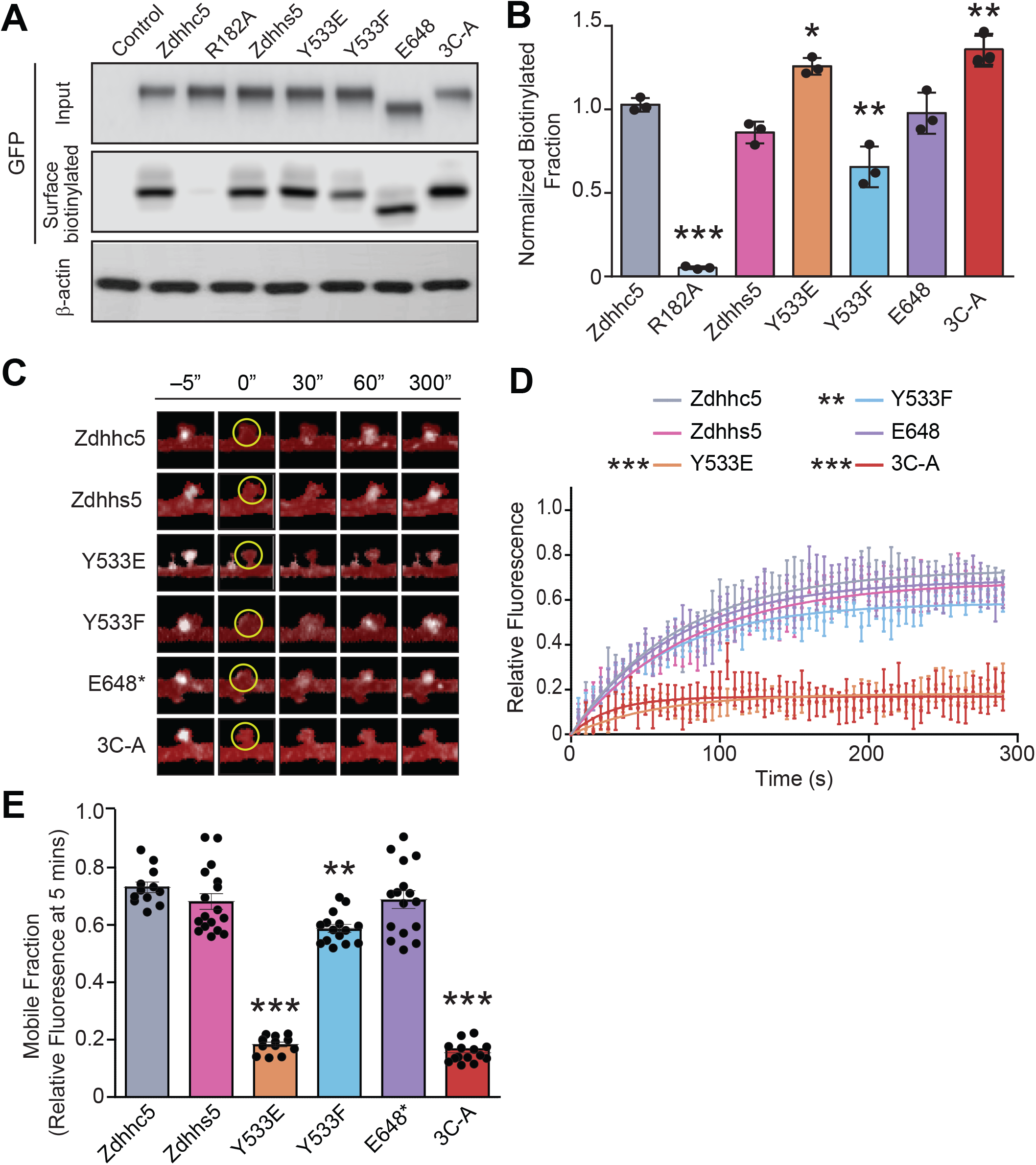
Zdhhc5 turnover is mediated by tyrosine 533 and C-terminal palmitoylation sites. (A-B) HEK293T cells transfected with sGFP-tagged Zdhhc5 constructs were biotinylated and lysates immunoprecipitated with neutravidin-coated beads to isolate all surface proteins and blots probed with anti-GFP to identify surface biotinylated Zdhhc5 mutants. The control represents a non-transfected sample and R182A a biotinylation control. The Y533E and 3C-A mutants were significantly more localized to the plasma membrane, while the Y533F mutant exhibited reduced surface localization. n = 3 blots, 3 separate cultures. *p < 0.05, **p < 0.01, ***p < 0.001; one-way ANOVA; Tukey’s post hoc; mean ± SEM. (C-E) Representative pseudocolored confocal images of neurons transfected with sGFP-tagged Zdhhc5 constructs following fluorescent recovery after photobleaching (FRAP) of a single dendritic spine. Spines were photobleached at 0 s within a 1µm2 ROI (yellow circle) (C). Y533E, Y533F, and 3C-A Zdhhc5 mutants exhibit significantly reduced mobile fractions (D, E). Statistical tests compare the plateau values from exponential fits ± SEM (D) or the relative fluorescence fraction within the ROI at the 5 min time point (E). n = 12-17 neurons, >3 cultures., **p < 0.01, ***p < 0.001; one-way ANOVA; Tukey’s post hoc; mean ± SEM.

We next expressed a C-terminal truncated Zdhhc5 identified in a patient with schizophrenia (Fromer et al., 2014), which results in a premature stop codon that deletes the last 68 amino acids of Zdhhc5 (Zdhhc5^R^ E648*), including the PDZ binding domain. Although surface expression of Zdhhc5^R^ E648* was similar to wildtype Zdhhc5 (Figure 3A, B), Zdhhc5^R^ E648* did not rescue the synaptic phenotype observed upon Zdhhc5 knockdown (Figure 2A-C). This deficit is likely due to reduced Zdhhc5 association with PDZ domain-containing proteins such as PSD-95, that are known to be critical for activity induced spine plasticity (Brigidi et al., 2015).

Our lab has previously demonstrated that phosphorylation of tyrosine 533 (Y533) immobilizes Zdhhc5 at the synaptic membrane by occluding Zdhhc5 binding to the endocytic protein, AP2µ (Brigidi et al., 2015). We observed a significant reduction in the surface localization of the Zdhhc5 phospho-dead mutant (Zdhhc5^R^ Y533F) (Figure 3A, B), which also failed to rescue Zdhhc5 knockdown-mediated decrease in excitatory synapse density (Figure. 2A-C). In contrast, while the phospho-mimetic Zdhhc5^R^ Y533E mutant was significantly *more* localized to the plasma membrane (Figure 3A, B), the density of excitatory synapses in cells expressing Zdhhc5^R^ Y533E was indistinguishable from cells expressing wildtype Zdhhc5.

Recent work has demonstrated that disrupting the palmitoylation of Zdhhc5 on cysteine residues 231, 238, and 245, impedes endocytosis and increases Zdhhc5 surface localization (Woodley & Collins, 2019; but see Ko et al., 2019). We therefore used a palmitoylation-defective, Zdhhc5^R^ 3C-A mutant to further explore the role of Zdhhc5 membrane localization in regulating synapse formation. As predicted, surface localization of Zdhhc5^R^ 3C-A was significantly increased when compared with wildtype Zdhhc5 (Figure 3A, B), yet no changes were observed in the density of excitatory synapses in cells expressing Zdhhc5^R^ 3C-A (Figure 2A, B).

Together, our data suggest that palmitoylation of protein(s) by Zdhhc5 at nascent synaptic membranes is required for the maturation or maintenance of synaptic connections. Decreased Zdhhc5 surface expression reduces excitatory synapse density; however, increased Zdhhc5 localization to the plasma membrane does not appear to be sufficient to promote synapse formation/maintenance.

We next investigated the mobility of Zdhhc5 mutants within postsynaptic spines using fluorescence recovery after photobleaching (FRAP). The mobile fraction of Zdhhs5 and Zdhhc5 E648* was unchanged compared to wildtype Zdhhc5 (Figure 3C-E). In contrast, we observed a decrease in the mobile fraction of Zdhhc5 Y533F, Y533E and Zdhhc5 3C-A (Figure 3C-E). Taken with the observation that surface localization of these three mutants was also significantly changed (Figure 3A, B), these results suggest an interplay between impaired mobility/trafficking and the compartmental accumulation of these mutants in either the plasma membrane (Zdhhc5-Y533E and −3C-A) or intracellular membranes (Zdhhc5-Y533F).

### *Zdhhc5 Regulates Excitatory Synapse Density* in vivo

To determine whether Zdhhc5 is similarly important for the formation and/or maintenance of excitatory synapses *in vivo*, we analyzed Zdhhc5 gene-trap (Zdhhc5-GT) mice (Li et al., 2010), in which Zdhhc5 protein is not detected (Figure. 4A). Using electron microscopy, we compared the density of synapses in the stratum radiatum of the hippocampus. Excitatory (asymmetric) and inhibitory (symmetric) synapses were identified based on their morphological characteristics (Gray, 1959; Figure 4B). To ensure accuracy, samples were also immunolabelled with PSD-95 to identify excitatory synapses (validated in Liu et al., 2016; Mills et al., 2017; Figure 4B). We observed a significant decrease in the density of PSD-95 positive, asymmetric excitatory synapses, but no change in the density of PSD-95 negative, symmetric inhibitory synapses in Zdhhc5-GT samples compared to controls (Figure 4C). No differences in active zone length were observed (Figure 4D).

**Figure 4.**
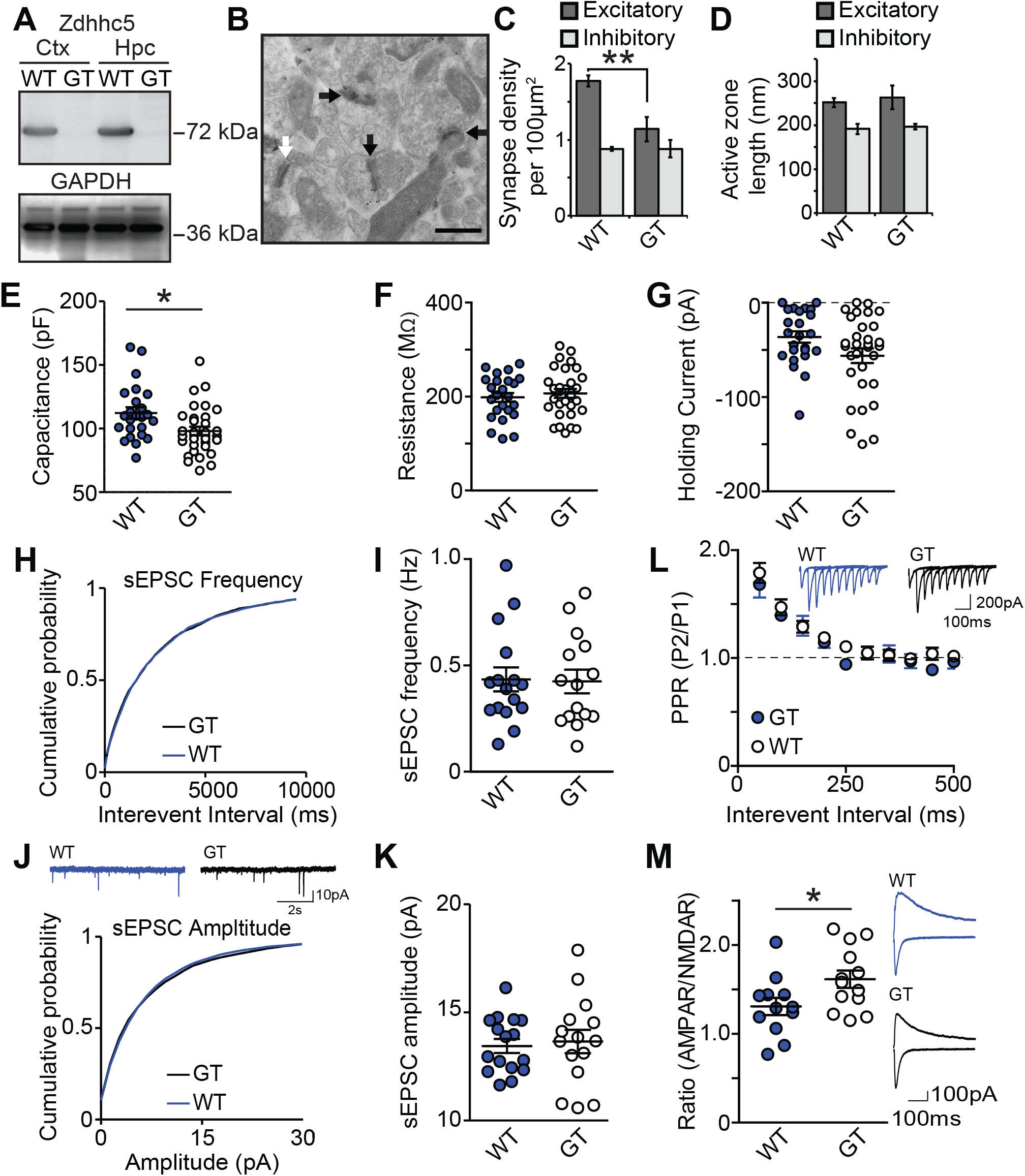
Zdhhc5 regulates excitatory synapse density in vivo. (A) Western blots of cortical (Ctx) and hippocampal (Hpc) lysates from Zdhhc5-GT (Gene Trapped) mice and age-matched littermate controls showing the absence of Zdhhc5 expression in Zdhhc5-GT mice. (B) Representative immunogold-electron microscopy images showing excitatory symmetric synapses (black arrows) and inhibitory asymmetric synapses (white arrows). Scale bar = 500 nm. (C) Zdhhc5-GT mice exhibit a significant reduction in the density of excitatory synapses, with no change in inhibitory synapses. n = 3 mice per genotype, two-way ANOVA; Bonferroni’s post hoc; significant interaction between genotype and synapse type, F(1, 8) = 8.753, p = 0.0182, **p significant interaction between genotype and synapse type, F(1, 8) = 8.753, p = 0.0182, **p<0.01. (D) There is no significant difference in the active zone between Zdhhc5-GT and WT littermates. (E) CA1 pyramidal neurons from acute Zdhhc5-GT brain sections exhibit lower capacitance than pyramidal neurons from WT mice (n = 24 WT neurons; 4 mice and 31 Zdhhc5-GT neurons; 5 mice; unpaired t-test, t(53) = 2.579, p=0.0127). (F-G) There is no significant difference between WT and Zdhhc5-GT mice in either resistance (F) or holding current (G). (H-J) There is no significant difference of sEPSC frequency (H, I), sEPSC amplitude (J, K), or paired-pulse ratio (L). (M) CA1 pyramidal neurons from Zdhhc5-GT brain slices exhibit a significantly higher AMPAR to NMDAR current ratio than WT slices. n = 12 WT neurons; 3 mice and 13 Zdhhc5-GT neurons; 3 mice; unpaired t-test, t(23) = 2.224, p=0.0363). All data is shown as mean ± SEM.

To determine whether the observed decrease in excitatory synapse density results in functional deficits, whole-cell voltage-clamp recordings were made in CA1 pyramidal neurons. The membrane properties revealed a lower capacitance in Zdhhc5-GT neurons compared to WT neurons (Figure 4E), with no change in membrane resistance or holding current (Figure 4F, G). As Zdhhc5 did not impact dendritic length or complexity *in vitro* (Figure 1A, B), and Zdhhc5-GT mice have a lower density of excitatory synapses (Figure 4C), this reduced capacitance may be due to fewer spines resulting in less total membrane.

Zdhhc5-GT and WT littermates revealed no difference in the frequency or the amplitude of sEPSCs (Figure 4H-K), despite our finding that Zdhhc5-GT mice have a decreased number of excitatory synapses. These observations could not be attributed to compensatory increases in pre-synaptic glutamate release in Zdhhc5-GT mice, as we found no significant difference in the paired pulse ratio (PPR) compared to WT littermates (Figure 4L). However, we observed a significantly higher AMPAR/NMDAR ratio in Zdhhc-GT mice (Figure 4M), without any changes in the amplitude of AMPAR sEPSCs (Figure 4J, K). These data may therefore reveal a selective decrease in the number of silent synapses (that contain NMDARs, but lack AMPARs (Isaac et al., 1995; Liao et al., 1995; Kerchner & Nicoll, 2008) in Zdhhc5-GT WT neurons. This is consistent with our observations that Zdhhc5 is required for spine maturation and stabilization, and would indicate that immature (silent) synapses might be eliminated before reaching maturity.

The findings of this study demonstrate that Zdhhc5, previously linked to learning and memory (Li et al., 2010) and schizophrenia (Schizophrenia Working Group of the Psychiatric Genomics, 2014), regulates excitatory synapse formation via the interplay of its palmitoylation activity, its C-terminal domain, and its surface localization. Two of the key neuropathological findings of schizophrenia are gray matter loss (reviewed in Vita et al., 2012), and changes in hippocampal dendritic spines (Steen et al., 2006; Kolomeets et al., 2005), while disruptions to excitatory/inhibitory balance underly a number of other neuropathologies. Our study contributes to the understanding of the role of Zdhhc5 in the hippocampus as a key component for learning and memory.

## Materials & Methods

### cDNA constructs

Control shRNA, Zdhhc5 shRNA, and HA-Zdhhc5 were kind gifts from R. Huganir (Johns Hopkins U., Baltimore). shRNA resistant HA-Zdhhc5 (Zdhhc5^R^) was generated as previously described (Brigidi et al., 2014). To make sGFP (Kremers et al., 2007)) Zdhhc5 constructs (sGFP-Zdhhc5-WT; -R182A; -E648; -Y533E; -Y533F and Zdhhs5), the coding sequences of mouse Zdhhc5 WT, R182A, E648, Y533E, Y533F and Zddhs5 were amplified from HA-Zdhhc5 constructs used previously (Brigidi et al., 2015) using the following primers: Fwd: 5’-CCGGCGAATTCTATGCCCGCAGAGTCTG-3’; Rev: 5’-GCCGGGGATCCTCACACAGAAATCTC-3’ for -WT, -R182A, -Y553E, -Y533F and Zdhhs5 or 5’-GCCGGGGATCCTCACTCTGAGACACCAGA-3’ for -E648. Fragments were then cut with EcoRI and BamHI and ligated into the multiple cloning site of the sGFP-C1 vector. To make sGFP-Zdhhc5-3A, the coding sequence of mouse Zdhhc5-WT was amplified with Gibson mutagenic primers to create 2 fragments incorporating the 3A mutations that were then joined using the NEB Gibson reaction mix (Fragment 1 primers: Fwd 5’-GATCTCGAGCTCAAGCTTCGAA-3’, Rev 5’-ACTGGCGAGGACACGGCTAACGTTGTTAGCGGCGCCATTGGTGAA-3’. Fragment 2 primers: Fwd 5’-ATGGCGCCGCTAACAACGTTAGCCGTGTCCTCGCCAGTTCTCCAGCA-3’, Rev 5’-TGATCAGTTATCTAGATCCGGTGG-3’). The joined Zdhhc5 3A fragment was then digested with EcoRI and BamHI and ligated into the sGFP-C1 vector.

### Cell Cultures

Hippocampi were isolated from embryonic day (E) 18 Sprague-Dawley rats as previously described (Xie et al., 2000) and plated at a density of 130 cells per mm^2^. Neurons were transfected with Lipofectamine 2000 (ThermoFisher, 11668019) at 10 days *in vitro* (DIV) following the manufacturer’s recommendations and used for experiments at 13 DIV. HEK293T cells were transfected using Lipofectamine 2000 (ThermoFisher, 11668019) according to the manufacturer’s recommendations. HEK293T cells were transfected at ∼70-80% confluency and incubated for 48 h before harvesting for biochemistry.

### Immunocytochemistry

Immunocytochemistry experiments were performed as previously reported (Sun and Bamji, 2011). Briefly, cultured neurons were fixed in 4% paraformaldehyde/4% sucrose, permeabilized with 0.1% Triton-X, and blocked in 10% goat serum for 1 h at room temperature. Primary antibodies were diluted in 1% goat serum and applied to neurons overnight at 4°C. Secondary antibodies were also diluted in 1% goat serum, and applied to neurons for 1 h at room temperature. Coverslips were mounted with Prolong Gold (ThermoFisher, P36930). Primary antibodies were as follows: PSD-95 (mouse monoclonal, IgG2a isotype, 1:500, abcam, ab2723), gephyrin (mouse monoclonal, IgG1 isotype, 1:500, Synaptic Systems, 147 011), VGLUT1 (guinea pig, 1:500, EMD Millipore, AB5905). Secondary antibodies were as follows: goat-anti-mouse IgG2a AlexaFluor 568 (Life Technologies, A21134), goat-anti-mouse IgG1 AlexaFluor 647 (Life Technologies, A21240), goat-anti-guinea pig AlexaFluor 633 (Life Technologies, A21105), goat anti-mouse AlexaFluor 568 (Life Technologies, A11019), goat anti-mouse AlexaFluor 633 (Life Technologies, A21050).

### Immunogold Electron Microscopy

Samples from mouse CA1 hippocampus were processed as described previously (Mills et al., 2017). Brains were sliced into 250 µm-thick sections by vibratome and pieces of CA1 hippocampus (<1 mm in all dimensions) were dissected from slices and cryoprotected in 30% glycerol overnight at 4°C. Samples were plunge-frozen in liquid ethane at −170°C in an EM cryopreparation chamber (Leica) and transferred to a 1.5% uranyl acetate solution in 100% methanol, kept at −90°C in a Leica EM AFS for 30 h. The temperature was gradually increased and samples infiltrated with HM-20 acrylic resin (Electron microscopy sciences, Hatfield, PA). Samples were set up in capsules containing pure resin and polymerized under UV light for 24h. Tissue sections were cut at 85 nm using a Diatome diamond knife and a Leica ultramicrotome. Sections were collected on 300-mesh, formvar coated nickel grids (Electron Microscopy Sciences).

Post-embedding immunostaining was performed on the EM grids as described previously (Mills et al., 2017). Grids were rinsed with distilled water and immersed in a bead of TTBS with 0.1% Triton-X, 0.1% sodium borohydride and 50 mM glycine. Nonspecific binding was blocked with 2% BSA in TTBS with 0.1% Triton-X. Primary antibodies PSD-95 (rabbit, Frontier Institute, Af628) were diluted in 2% BSA in TTBS with 0.1% Triton-X. Grids were immersed in 15 µl beads of diluted primary antibody overnight at room temperature in a humidified chamber. The next day, grids were rinsed in TTBS with 0.1% Triton-X. Secondary antibodies were diluted in 2% BSA in TTBS with 0.1% Triton-X and 0.05% polyethylene glycol (PEG). Girds were immersed in 15µl beads of secondary antibody (Electron microscopy sciences, goat-anti-rabbit 15 nm, cat. no. 25112) for 1.5 h. Grids were rinsed in TTBS with 0.1% Triton-X, in Milli-Q H2O and dried. Grids were then lightly counterstained with 2% uranyl acetate and Reynold’s lead citrate. Images were collected at 23,000x magnification on a Tecnai G2 Spirit transmission electron microscope (FEI Company, Eindhoven, the Netherlands). To quantify excitatory and inhibitory synapse density in the CA1 hippocampus, the number of PSD-95 positive and negative synapses was quantified within a 2500 µm^2^ region of the hippocampus. All images were acquired and analyzed blind to the genotype of each mouse.

### Fluorescence Recovery After Photobleaching (FRAP)

sGFP-Zdhhc5 and sGFP-Zdhhc5 mutant construct puncta localized to dendritic spines (identified using mCherry cell fill) within 100 µm of the cell body were imaged every second for 5 minutes after photobleaching using a Zeiss LSM780 confocal microscope. An approximate 1µm diameter circular region of interest was used to photobleach the Zdhhc5, and a 1µm diameter circular region of interest was placed both on the cell body for a reference and a region of background for background substraction. Analysis was carried out using easyFRAP (Rapsomaniki et al., 2012) and data exported to Prism Software (GraphPad) for data visualization, analysis and plateau value generation.

### Immunoblot (Western) Assay

Western blotting was performed as previously described (Brigidi et al., 2014; Sun and Bamji, 2011). HEK293T cells were homogenized in an ice-cold lysis buffer containing 1% IGEPAL CA-630 (Sigma), 50 mM Tris-HCl, pH7.5, 150 mM NaCl and 10% glycerol, supplemented with phenylmethanesulfonyl fluoride solution and a protease inhibitor cocktail with ethylenediaminetetraacetic acid (Roche). Brain tissue was homogenized in ice-cold RIPA buffer (ThermoFisher, CAT# 89900). Proteins were cleared by centrifugation at 14,000g for 30 min at 4°C. Proteins were separated by SDS-PAGE and probed with antibodies against zDHHC5 (1:1,000; Sigma Prestige, HPA014670), HA (1:1,000; Sigma, H9658), and GAPDH (1:1,000; abcam, ab9484). Bands were visualized using enhanced chemiluminescence (Pierce Biotechnology) on a C-DiGit Chemiluminescence Western Blot Scanner (LI-COR).

### Biotinylation Assay

Biotinylation experiments were performed as previously described (Shimell et al., 2019). Briefly, neurons in 10cm dishes were nucleofected with indicated constructs and experiments were carried out at 13 DIV. Neurons were washed with ice cold PBS-CM (0.1 mM CaCl2 and 1 mM MgCl2 in 1X PBS, pH 8) and incubated for 30 min with 0.5 mg/mL NHS-SS-Biotin in ice cod PBS-CM at 4°C with gentle rocking. After incubation cells were washed once with PBS-CM and the unbound biotin quenched via two 8 min incubations with quenching buffer (20 mM glycine in PBS-CM). Lysis was performed using mechanical scraping in lysis buffer (1% IGEPAL-CA630 and 1mMPMSF with Roche complete protease inhibitor tablet) and subsequently spun down at 500xg for 5 minutes at 4°C. Samples were vortexed, run through a 26 1/2-gauge syringe 3 times, and nutated at 4°C for 30 min. After nutation, samples were spun down at 16100xg for 30 minutes at 4°C to clear the lysate. The cell lysate was then quantified for protein using a BCA Assay Kit (Thermo Fisher Scientific) as per the manufacturer’s instructions. 10 mg of each whole cell lysate was then combined with SDS-sample buffer (50 mM Tris-HCl, 2% SDS, 10% glycerol, 14.5 mM EDTA and 0.02% bromophenol blue with 1% β-mercaptoethanol), boiled for 5 min at 95°C and stored at −20°C as the input sample. 100-200 mg of the remaining protein sample was added to a 50 mL 50% slurry of Neutravidin-conjugated agarose beads (Thermo Fisher Scientific) that was pre-washed 3X in lysis buffer. Each sample was then brought to a total volume of 500 mL with lysis buffer and nutated at 4°C overnight. The following day beads were pelleted and washed 7X using centrifugation (500xg for 3 min). Elution of the beads was performed using 40 mL of SDS-sample buffer with 100mMDTT. Samples were boiled at 90°Cfor 5 min and then run on a western blot with the whole cell lysates.

### Calcium Imaging with GCaMP6f

Cultured hippocampal neurons were transfected on 12 DIV with pCAG-GCaMP6f (0.2 ug), and either control shRNA (1 ug); Zdhhc5 shRNA (1ug); or Zdhhc5 shRNA (1 ug) and WT HA-Zdhhc5^R^ (1 ug). At 15 DIV, neurons were incubated at 20°C in an artificial CSF (aCSF) imaging media containing the following in mM: 135 NaCl, 5 KCl, 25 HEPES, 10 glucose, 3 CaCl_2_, and 0.001 TTX, pH 7.4. Images were acquired using a Zeiss LSM 880 AxioObserver Airyscan microscope with a Plan-Apochromat 63x/1.4 Oil DIC M27 objective and a 0.71 μs dwell time (488 nm laser) using AiryScan Fast mode. Single z-plane images of the proximal dendritic arbor covering a 133.5 × 133.5 μm field of view were acquired at 3.8 Hz for 2 minutes.

To measure the number and amplitude of Ca^2+^ events, the time-lapse image was processed using the “Delta F up” plugin from the ImageJ Cookbook T-functions application and then maximum intensity projected to create a binary mask of Ca^2+^ event locations, which were then converted into regions of interest (ROIs). Any events that occurred in presumed axons or the soma were removed. The mean GCaMP6f fluorescence within each ROI was measured. A baseline of 10 frames was established for each ROI and used to calculate the ΔF/F_0_. Ca^2+^ events were counted that surpassed a threshold of a 200% increase in fluorescence over baseline. The mean number of events per spine was calculated as the total number of active spines divided by the total number of Ca^2+^ events. Ca^2+^ events per μm were calculated as the total number of active spines (classified as spines with ≥ 1 Ca^2+^ event) divided by total dendrite length, which was measured using the “NeuronJ” plugin in ImageJ (Meijering et al., 2004).

### Slice Preparation and Electrophysiology

Male 2-month-old zDHHC5-gt (Li et al., 2010) and wildtype control mice were deeply anesthetized with isofluorane vapor, decapitated and the brain rapidly removed. Acute sagittal brain slices (300µm) containing medial dorsal hippocampus were cut on a vibratome (Leica VT1000) in ice-cold artificial cerebrospinal fluid (aCSF; with 0.5 mM CaCl_2_ and 2.5 mM MgCl_2_) equilibrated with 95% O2/5% CO_2_. Slices were transferred to a holding chamber with aCSF at 35°C containing (in mM): 125 NaCl, 2.5 KCl, 25 NaHCO_3_, 1.25 NaH_2_PO_4_, 1 MgCl_2_, 2 CaCl_2_, 10 glucose, pH 7.3–7.4, 305–310 mosmol L^−1^ for 45 minutes then maintained at room temperature. In the recording chamber, slices were equilibrated for 10 minutes while being continuously superfused at room temperature with oxygenated aCSF at 1-2 ml/min containing picrotoxin (50 μM, Tocris Bioscience, MO, USA) to block GABA_A_ receptor-mediated inhibitory responses.

Pipettes (3-5MΩ) were made from borosilicate glass capillaries on a Narishigi micropipette puller (Narishige International, East Meadow, New York). Whole cell patch-clamp recording was performed with a multiclamp-900 amplifier and pClamp 10 software (Axon instruments, CA, USA) digitized at 20 kHz and filtered at 10 kHz (sEPSC were filtered at 1 kHz with a detection threshold of −8 pA). CA1 pyramidal neurons were maintained voltage-clamped at - 70mV for sEPSC, PPR and AMPAR/NMDAR ratio (A/N ratio, except that the voltage was switched to +40mV to measure NMDAR current) experiments with internal solution containing in mM: 120 Cesium Methane-Sulfonate (CH_3_O_3_SCs), 5 NaCl, 1.1 EGTA, 4 MgATP, 0.3 NaGTP, 5 QX-314 Cl, 10 tetraethylammonium chloride (TEA), 10 HEPES, pH 7.25, osmolarity 290 mOsm. Series resistance was <20mΩ and uncompensated. To evoke synaptic currents, stimuli (100μsec duration) were delivered with a glass electrode (2-3MΩ) filled with aCSF placed <300 µm from the recorded cell to stimulate the Schaffer collateral pathway. Paired - pulse ratios (PPR) were calculated by dividing the amplitude of the second EPSC by the amplitude of the first, and increasing the interpulse interval by 50ms to a maximum of 500ms and repeated twice for each cell. For A/N ratio, the AMPAR component was the average peak amplitude of 4 evoked EPSC at −70mV. The NMDAR component was the average of 4 EPSC evoked at the same stimulation intensity and clamped at +40mV measured 75ms after stimulation to eliminate the possibility of contamination by AMPAR.

### Image Acquisition, Analysis, and Quantification

Confocal images were obtained on either an Olympus Fluoview FV1000 inverted laser scanning confocal microscope or a Zeiss 880 laser scanning confocal microscope using a 60X or 63X imaging objective, respectively. For synapse density analyses, confocal images were subjectively thresholded using ImageJ software (observer blind to condition). Puncta were identified as a fluorescence cluster with an area between 0.05 and 3 μm^2^. Puncta area and the integrated density (the product of the area and the mean gray value) were then determined using ImageJ. An ImageJ colocalization plugin was used to assess the colocalization between VGLUT1 and PSD-95 puncta (http://rsb.info.nih.gov/ij/plugins/colocalization.html). Points of colocalization were defined as regions of >4 pixels in size with a >50 intensity ratio between the two channels. The length of each dendrite was measured using the “NeuronJ” ImageJ plugin (Meijering et al., 2004), and puncta density was calculated by computing the number of puncta in each image divided by the length of dendrite in each image. Sholl analysis was performed using an ImageJ plugin (Ferreira et al., 2014). Reconstruction and analysis of dendritic spine type was measured using NeuronStudio computational software (Rodriguez et al., 2008).

### Statistical Analysis

All data values are expressed as the mean ± SEM. For imaging experiments, the n numbers shown refer to the number of cells used per condition over 3 separate cultures, with the exception of Figure 3A-B (FRAP), where *N* refers to the number of spines and is specified within figure legends. All electrophysiological data were analyzed using Clampfit 10.4 and Graphpad Prism 5 and are presented as the mean +/- SEM. of n = number of neurons from a minimum of 3 animals per genotype. Statistical significance was determined by Student’s *t* test, one-way or two-way ANOVA with Bonferroni’s test *post hoc* using Prism where indicated. Statistical significance was assumed when *p* < 0.05. All figures were generated using Photoshop CS6 and/or Illustrator CS6 software (Adobe Systems, Inc.).

## ACKNOWLEDGEMENTS

This work was supported by grants from CIHR (F18-00650, S.X.B; FDN-143210, LAR).

---

**Supplementary Figure 1.**
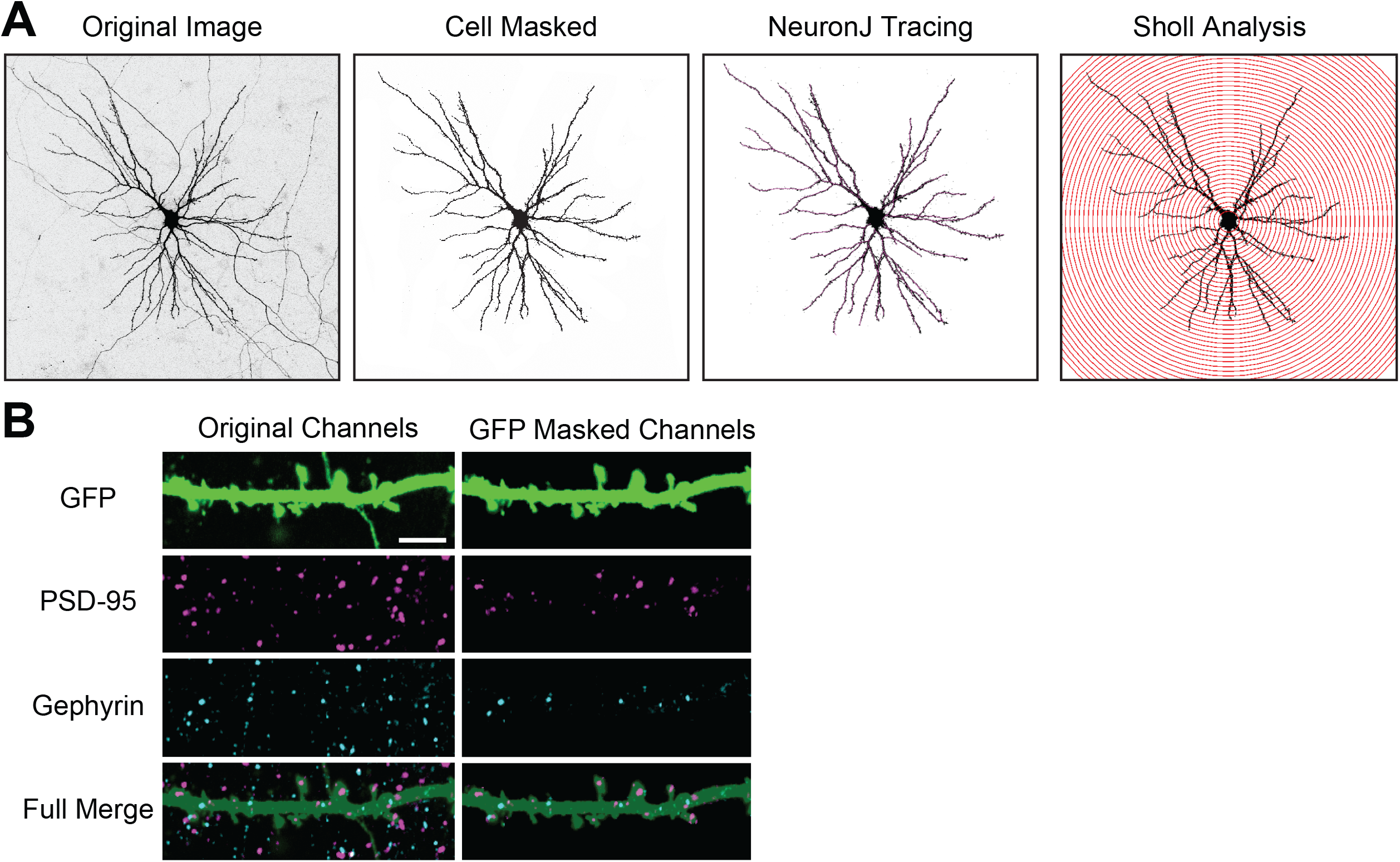
GFP masking for neurons tracing and quantification of synapse density. (A) Representative eGFP transfected neuron. The original image is manually masked to remove background and axons and then traced using the NeuronJ plugin for ImageJ. (B) Representative eGFP transfected neurons immunostained with the excitatory marker, PSD-95, and the inhibitory marker, gephyrin. The right column shows unmasked images, while the left column is masked using the eGFP channel.

